# Genetic Analysis of Human RNA Binding Motif Protein 48 (RBM48) Reveals an Essential Role in U12-Type Intron Splicing

**DOI:** 10.1101/2020.07.18.209528

**Authors:** Amy E. Siebert, Jacob Corll, J. Paige Gronevelt, Laurel Levine, Linzi M. Hobbs, Catalina Kenney, Ruth Davenport, A. Mark Settles, W. Brad Barbazuk, Randal J. Westrick, Gerard J. Madlambayan, Shailesh Lal

## Abstract

U12-type or minor introns are found in most multicellular eukaryotes and constitute ∼0.5% of all introns in species with a minor spliceosome. Although the biological significance for evolutionary conservation of U12-type introns is debated, mutations disrupting U12 splicing cause developmental defects in both plants and animals. In human hematopoietic stem cells, U12 splicing defects disrupt proper differentiation of myeloid lineages and are associated with myelodysplastic syndrome (MDS), predisposing individuals to acute myeloid leukemia. Mutants in the maize ortholog of RNA Binding Motif Protein48 (RBM48) have aberrant U12-type intron splicing. Human RBM48 was recently purified biochemically as part of the minor spliceosome and shown to recognize the 5’ end of the U6atac snRNA. In this report, we use CRISPR/Cas9-mediated ablation of *RBM48* in human K-562 cells to show the genetic function of RBM48. RNA-seq analysis comparing wild-type and mutant K-562 genotypes found that 48% of minor intron containing genes (MIGs) have significant U12-type intron retention in *RBM48* mutants. Comparing these results to maize *rbm48* mutants defined a subset of MIGs disrupted in both species. Mutations in the majority of these orthologous MIGs have been reported to cause developmental defects in both plants and animals. Our results provide genetic evidence that the primary defect of human *RBM48* mutants is aberrant U12-type intron splicing, while a comparison of human and maize RNA-seq data identifies candidate genes likely to mediate mutant phenotypes of U12-type splicing defects.

## Introduction

Accurate splice site recognition and precise removal of introns during pre-mRNA processing is fundamental for gene expression in eukaryotes^1-3^. The vast majority of introns are spliced by the major spliceosome and are categorized as U2-type (or major) introns^4, 5^. A second group of introns are spliced by the minor spliceosome and are categorized as U12-type (or minor) introns^6^. U12-type introns constitute ∼0.5% of all introns in species with a minor spliceosome^6^. In contrast to U2-type, U12-type introns have highly conserved 5’ splice site and branch point sequences, tolerate distinct 5’ and 3’ terminal dinucleotides, and lack 3’ polypyrimidine tracts^77, 88^. Both U12-type introns and U2-type introns exist side-by-side within Minor Intron-containing Genes (MIGs)^6^. Most of the 758 MIGs identified in humans contain only a single U12-type intron^9^.

Defects in pre-mRNA splicing by cis-or trans-acting mutations have been linked to approximately 60% of human diseases^10-12^. While many mutations impact splicing of U2-type introns^13^, anomalous splicing of U12-type introns can cause developmental defects in both plants and animals^14-19^. Mutations in minor spliceosome factors impact normal splicing of a subset of MIGs in both humans and maize that result in cell differentiation defects^20-22^. For example, aberrant differentiation and proliferation of myeloid precursors associated with myelodysplastic syndrome (MDS) can be caused by mutations in the spliceosomal gene *ZRSR2*, which impairs splicing of U12-type introns leading to excess intron retention^22^. MDS patients with *ZRSR2* mutations display peripheral cytopenia and accumulate abnormal myeloid cells due to defective hematopoiesis of myeloid precursors in the bone marrow, resulting in an increased risk to acute myeloid leukemia (AML)^23^. In maize, the corresponding *ZRSR2* ortholog is disrupted by the hypomorphic *rough endosperm3* (*rgh3*) allele which results in aberrant endosperm differentiation and proliferation as well as impaired splicing and retention of U12-type introns^21^. A comparison of MIGs affected in the human *ZRSR2* and maize *rgh3* mutants identified conserved cellular and molecular pathways associated with cell proliferation and expression of terminal cell fates^21^.

Major and minor spliceosomes are complex molecular machines displaying remarkable similarity^6^. Many of the core and auxiliary protein components are postulated to be shared between the two complexes. To date, only 12 have been identified to be specific to the minor spliceosome^66, 24^. One of these U12-specific splicing factors, RNA Binding Motif Protein 48 (RBM48) is co-conserved in eukaryotes with ZRSR2 and shown to be required for efficient splicing of U12-type introns in maize^25^. Maize *rbm48* mutants share a stark phenotypic similarity with *rgh3* mutants in endosperm developmental defects, as both loci exhibit increased cell proliferation and decreased cell differentiation^25^. Maize RBM48 interacts with core U2 Auxiliary Factor (U2AF), Armadillo Repeat-Containing Protein 7 (ARMC7), and RGH3, suggesting the components of both major and minor spliceosomal complexes interact during pre-mRNA processing. A recent structural analysis of a purified human minor spliceosome complex confirmed the interaction of RBM48 with ARMC7. This complex in turn binds to the 5’ cap of U6atac, potentially facilitating U12 splicing by stabilizing the intronic sequence in addition to the minor spliceosome itself^24^. However, there is no direct genetic evidence demonstrating a U12 splicing role for RBM48 in human cells.

Herein, we generated a CRISPR/Cas9 mediated functional knockout of *RBM48* in human K-562 chronic myeloid leukemia cells. Using comprehensive transcriptome profiling, we demonstrate that RBM48 is required for efficient splicing of U12-type introns indicating conservation of function between plants and humans. Comparative analyses identified a subset of conserved MIGs that are affected in both maize and human RBM48 knockout mutants. These data identify biological processes impacting cell proliferation and development that have maintained common post-transcriptional RNA processing mechanisms since the divergence of plants and animals.

## Results

### Targeting human *RBM48* for Cas9-mediated cleavage in K-562 cells

The CRISPR/Cas9 genome editing system was implemented to generate a functional knockout of *RBM48* in K-562 cells. Two single-guide RNAs (sgRNA) were designed to target the N-terminal (sgRNA#1) and C-terminal (sgRNA#2) regions of the RNA Recognition Motif (RRM) domain within the *RBM48* genomic locus. The position, sequence, and approximate cleavage site of each sgRNA is displayed schematically in Figure 1A. To determine the efficacy of Cas9-mediated DNA cleavage at each targeting site, genomic DNA (gDNA) was analyzed by the Surveyor nuclease assay from cell populations at harvest following puromycin selection (day 0), as well as 10 and 16 days post-selection (Fig. 1B). Insertion-deletion (indel) polymorphisms generated within the *RBM48* genomic locus were detected on day 0 targeted cell populations by heteroduplex cleavage fragments of 309 and 272 base pair (bp) for sgRNA#1- and 332 and 266 bp in sgRNA#2. No detectable fragments were observed in digested vector control (VC) amplicons. Indels were detected in each targeted population at 10 days post-selection. By day 16, digested fragments were undetectable from sgRNA#1-targeted DNA. By contrast, cleavage fragments from DNA targeted by sgRNA#2 were stable and appeared to intensify at 16 days post selection.

**Figure 1.**
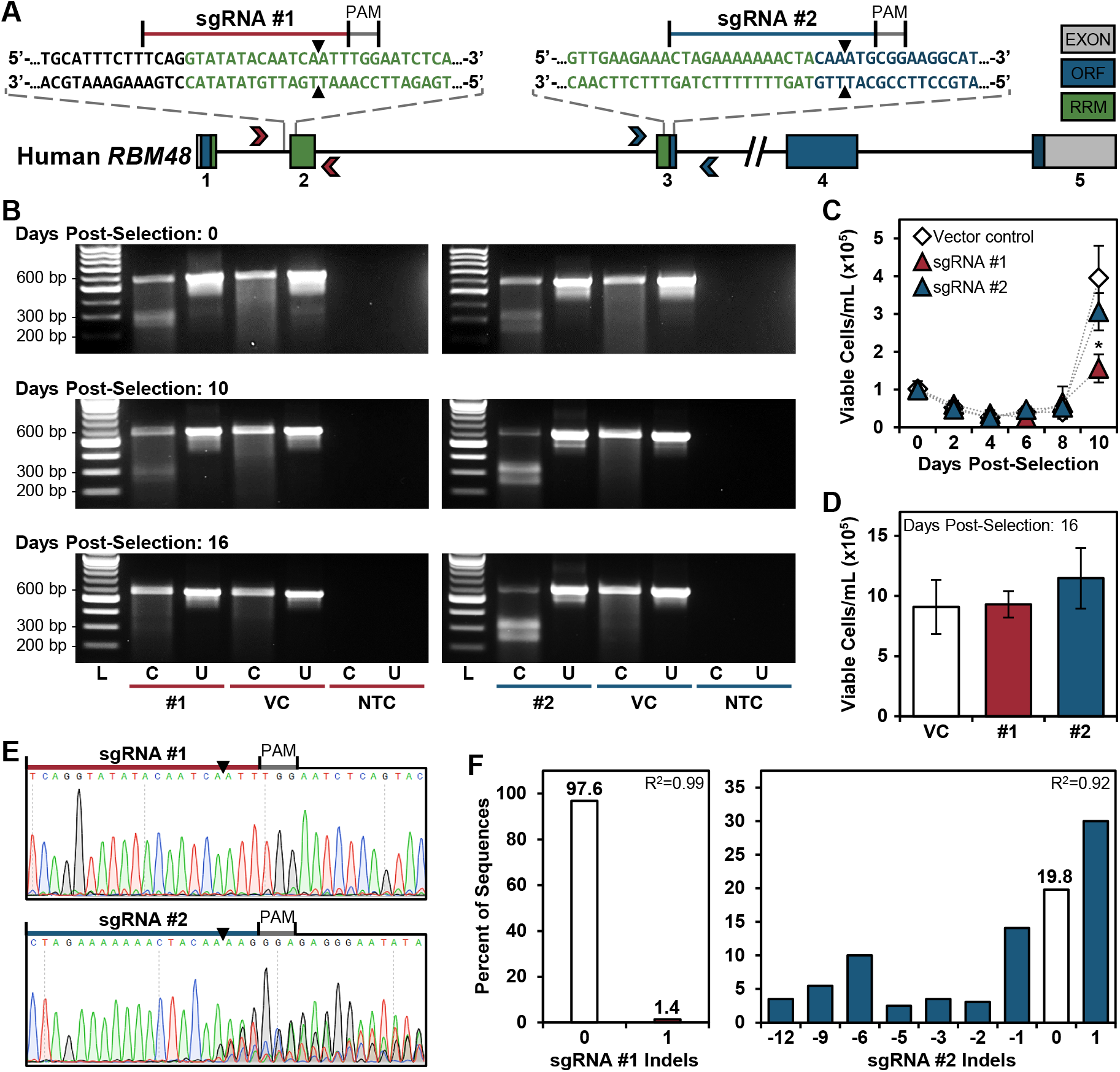
Functional CRISPR/Cas9-mediated knockout of *RBM48* in K-562 cells. **(A)** Schematic of human *RBM48* gene (NM_032120.4) structure displaying the design and position of the sgRNAs used for Cas9 targeting. Gray, green, and blue boxes indicate exons with the exon number indicated below each box. The open reading frame (ORF) is blue, and the RNA Recognition Motif (RRM) is green. Chevron arrowheads indicate position and direction of PCR primers used in Figure 1B, E and F for sgRNA#1 (red) and #2 (blue) and are listed in Table S3. The target site of each sgRNA is expanded above the structure. Black, blue, and green text are intronic, ORF and RRM domain sequences, respectively. Black arrowheads mark the predicted cleavage site of Cas9. PAM is the protospacer adjacent motif. **(B)** Surveyor Nuclease Assay. Heterogeneous K-562 cell populations transfected with VC or sgRNA#1 (left panel set, red underline) or sgRNA#2 (right panel set, blue underline), were amplified and digested with Surveyor nuclease to determine indel formation within transfected cell populations at 0, 10 and 16 days post-selection. The expected sizes for uncut (U) amplicons spanning the regions targeted by sgRNA#1 and #2 were 581 and 599 bp, respectively. C = cut with surveyor nuclease. NTC = no template control. The molecular weight standards are indicated on the left. **(C)** and **(D)** Cell viability assays of transfected K-562 cells. Viable cell counts monitored every 48 h from 72 h post-puromycin selection (day 0) through day 10 are shown in (C). \**q*<0.05. Viable cell counts on day 16 are shown in (D). **(E)** Sanger sequencing chromatograms of the amplified region flanking the Cas9 cleavage site targeted by sgRNA#1 (top panel) and sgRNA#2 (bottom panel) at day 16 post-selection. **(F)** TIDE analysis of the chromatograms displayed in Figure 1E. Genomic DNA amplified from VC cell populations was used as non-targeted K-562 input sequence for the analysis.

Growth of the transfected cell populations was monitored over the 16-day post-selection time interval (Fig. 1C and D). Over the first 8 days, cell growth was reduced in all 3 cohorts as a result of the transfection procedure (Fig. 1C). While the VC and sgRNA#2 cells began to recover at 10 days post-selection, proliferation lagged in the sgRNA#1-targeted K-562 cell population as compared to VC cells (*q*=0.002). By day 16 however, the sgRNA#1 cells regained vigor with a comparable number of viable cells from both Cas9-targeted and VC populations (Fig. 1D).

The indels generated within the two *RBM48*-targeted cell populations were further investigated by Sanger sequencing and Tracking of Indels by DEcomposition (TIDE) analysis on day 16 post-selection cells. Sequencing chromatograms flanking the protospacer adjacent motif (PAM) site of sgRNA#1 (Fig. 1E; upper panel) and subsequent TIDE analysis (Fig. 1F; left), showed that the sgRNA#1-targeted cells were a nearly homogenous population predominantly harboring wild-type *RBM48* by day 16 post-selection. By contrast, multiple sequence trace peaks immediately following the sgRNA#2 Cas9 cleavage site were observed (Fig. 1E; lower panel). The frequency of wild-type *RBM48* sequences within the heterogeneous sgRNA#2 cell population by TIDE analysis was only 19.8%, while the total Cas9 cleavage efficiency was approximately 72% with nearly 30% comprising a 1 bp insertion (Fig. 1F; right). From these observations, we conclude that sgRNA#1 was inefficient in generating stable targeted-indels, which led to the proliferation of unmodified *RBM48* wild-type cells as the dominant population. However, the sgRNA#2 targeting site was highly efficient at generating stable indels without apparent growth defects, enabling us to readily propagate this cell population. Therefore, the VC and sgRNA#2-targeted cell populations were established and maintained as independent K-562 cell sublines, hereafter referred as VC and *RBM48* KO, respectively.

### Loss of full length RBM48 in the *RBM48* KO subline

Loss of *rbm48* in maize impairs splicing of U12-type introns^25^. To elucidate whether human RBM48 has a conserved role in U12-dependent splicing, we performed mRNA-seq analysis on six unique passages of both *RBM48* KO and corresponding VC populations. Prior to analyzing the mRNA-seq data for splicing defects, *RBM48* expression was compared between the heterogeneous KO and VC populations to determine the impact of CRISPR/Cas9 mutagenesis. In agreement with the sgRNA#2 TIDE analysis in Figure 1F, a disparity in summed read depth was clearly visible within the targeting region of *RBM48* exon 3 in the KO population in comparison to VC (Fig. 2A). However, RT-qPCR assays for expression of all three *RBM48* transcripts (Fig. 2B) showed no significant differences between the KO and VC populations over six passages indicating that Cas9-targeting did not affect overall transcription of the *RBM48* gene. Through further analysis, 12 indel-harboring transcript sequences were identified that were maintained within all six passages of the *RBM48* KO subline. Figure 2C displays the indel positions and their predicted translational consequences in relation to NM_032120.4 and NP_115496.2, respectively. Within the heterogeneous KO subline, we found evidence for two mutant transcripts resulting in loss of a single amino acid and ten mutant transcripts resulting in frameshifts and truncation of the C-terminal region of the RBM48 protein product in all three coding variants.

**Figure 2.**
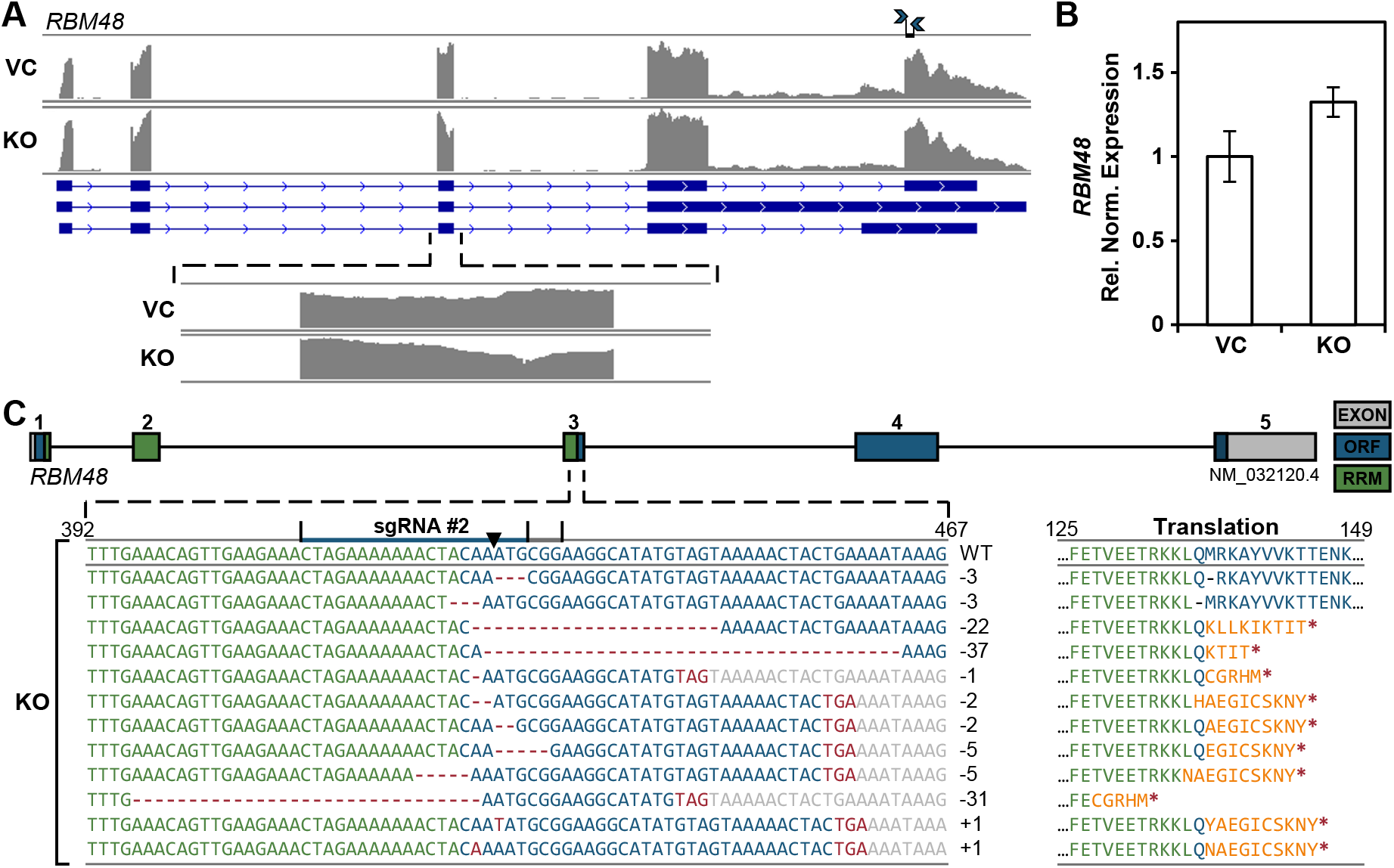
*RBM48* transcript expression from the KO cell population. **(A)** RNA-seq read depth comparison of *RBM48* expression between summed VC and KO libraries. Blue arrowheads mark the position of primers used for RT-qPCR in (B). The sequence of the primers is listed in Table S5. The expanded bottom panel displays the region spanning the sgRNA#2 target site in exon 3 of the *RBM48* sequence. **(B)** RT-qPCR of relative *RBM48* transcript expression between *RBM48* KO and VC cells are displayed as the mean ± SEM from six serial cell passages (n = 6 per cell subline) and normalized to reference genes *HPRT1, IPO8*, and *PGK1*. **(C)** *RBM48* KO expressed transcripts determined from RNA-seq. The left panel displays the sequences from the *RBM48* sgRNA#2-targeting region spanning positions 392-467 of the NM_032120.4 transcript variant. The number of nucleotide insertions/deletions (indels) within each sequence are indicated. The predicted translational product of each transcript starting amino acid position 125 of the NP_115496.2 protein accession is shown on the right. Orange text designates changes in protein sequence. Red asterisks mark premature stop codons.

We then generated a third K-562 cell subline that carried a Cas9-mediated C-terminal Myc epitope tag (*RBM48-Myc*; described in Figure S1) as a tool to investigate protein truncation and loss of full-length expression by sgRNA#2 targeting. The *RBM48* C-terminus was selected for epitope tagging to avoid a potentially lethal synthetic interaction due to overlap between the *RBM48* start codon site and promoter regulatory regions of the proximal *PEX1* gene (ENSEMBL Regulatory Feature: ENSR00000215089). CRISPR/Cas9 targeting in the *RBM48-Myc* subline using sgRNA#2 (KO) resulted in a significant reduction in tagged RBM48-Myc protein levels in comparison to VC (Fig. 3A and B; *p*=0.0051) This finding provides evidence that indels generated by sgRNA#2 targeting of *RBM48* exon 3 could result in at least C-terminal protein truncation and reduced expression of the full length RBM48 protein product.

**Figure 3.**
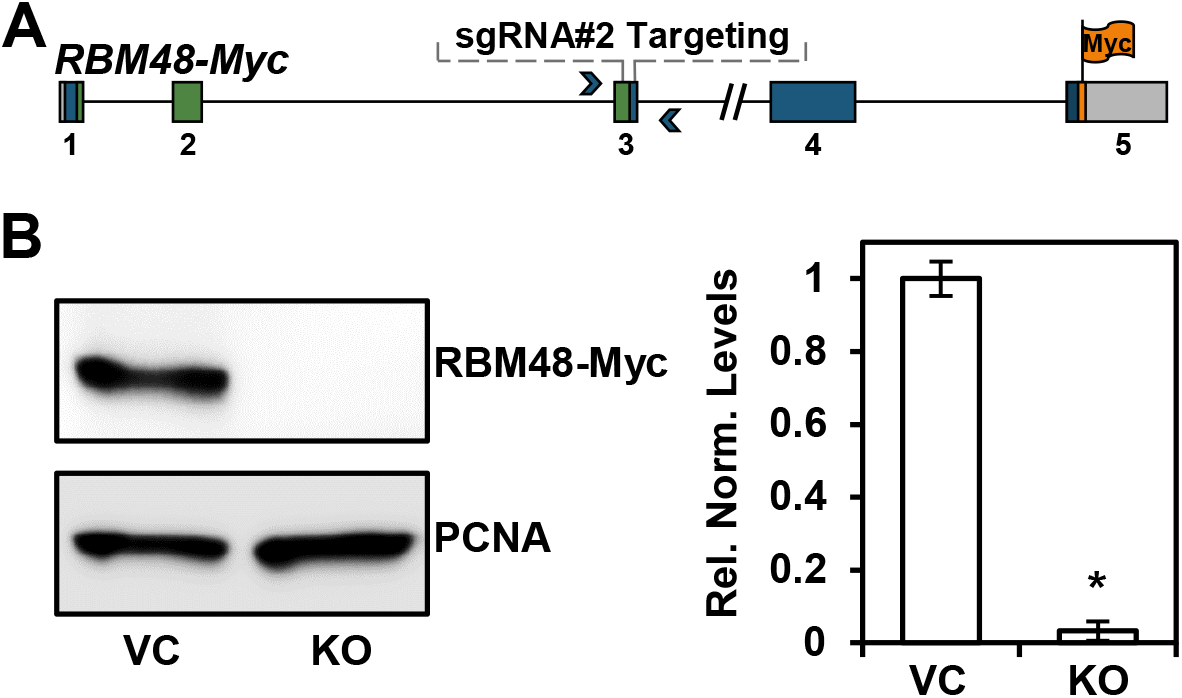
Ablation of RBM48-Myc protein levels by sgRNA#2-targeting of Cas9. **(A)** Schematic of *RBM48*-*Myc* displaying the sgRNA#2 targeting site. **(B)** Western blot (left panel) and densitometric analysis (right panel) of RBM48-Myc protein levels between KO and VC cell populations. The relative intensities of RBM48-Myc are displayed as the mean ± SEM from three independent cell passages (n = 3) and normalized to PCNA levels. \**p*<0.001 by Student’s t-test.

### Disruption of U12-dependent splicing in *RBM48* KO cells

To assess whether the *RBM48* RNA processing role is conserved in maize and humans, transcriptome profiles of the *RBM48* KO and VC populations were analyzed for intron splicing defects. Intronic and flanking exon-exon junction read counts for each individual U2- and U12-type intron expressed in the K-562 cell sublines were assessed for significance using Fisher’s Exact Test (Dataset S1). A minimum sum of 10 total exon-exon junction and intron reads with an intron read density > 0 in both VC and KO populations were required to test an intron. Read count for each individual sample is provided in Dataset S2. Of the 529 U12-type introns tested, 253 (47.8%) were significantly retained in the *RBM48* KO population in comparison to VC (3.7%; *q* ≤ 0.05). Gene Ontology (GO) term enrichment analysis identified cellular localization as the most significantly enriched term (*q* ≤ 0.05) among MIGs with U12-type intron retention (Table S1).

Four affected U12-type introns in the *DIAPH1, MAPK1, MAPK3*, and *TXNRD2* genes were selected for validation by RT-qPCR. Based on our mRNA-seq data, each of these genes exhibited significantly increased read depth (*q* ≤ 0.05) in the U12-type intronic regions in *RBM48* KO populations compared to VC (Fig. 4). We designed primers to detect both U12-type intron retention and total transcript levels by RT-qPCR for each of the selected MIGs (Fig. 4). No significant differences were found in the total transcript expression levels between the *RBM48* KO and VC populations. By contrast, expression levels of all four U12-type introns were significantly increased, suggesting intron retention in these transcripts. These findings indicate that in humans, RBM48 plays a role in post-transcriptional RNA processing and U12-dependent splicing, but that U12-type intron retention does not necessarily reduce the transcript levels of all genes where the U12-type introns are retained.

**Figure 4.**
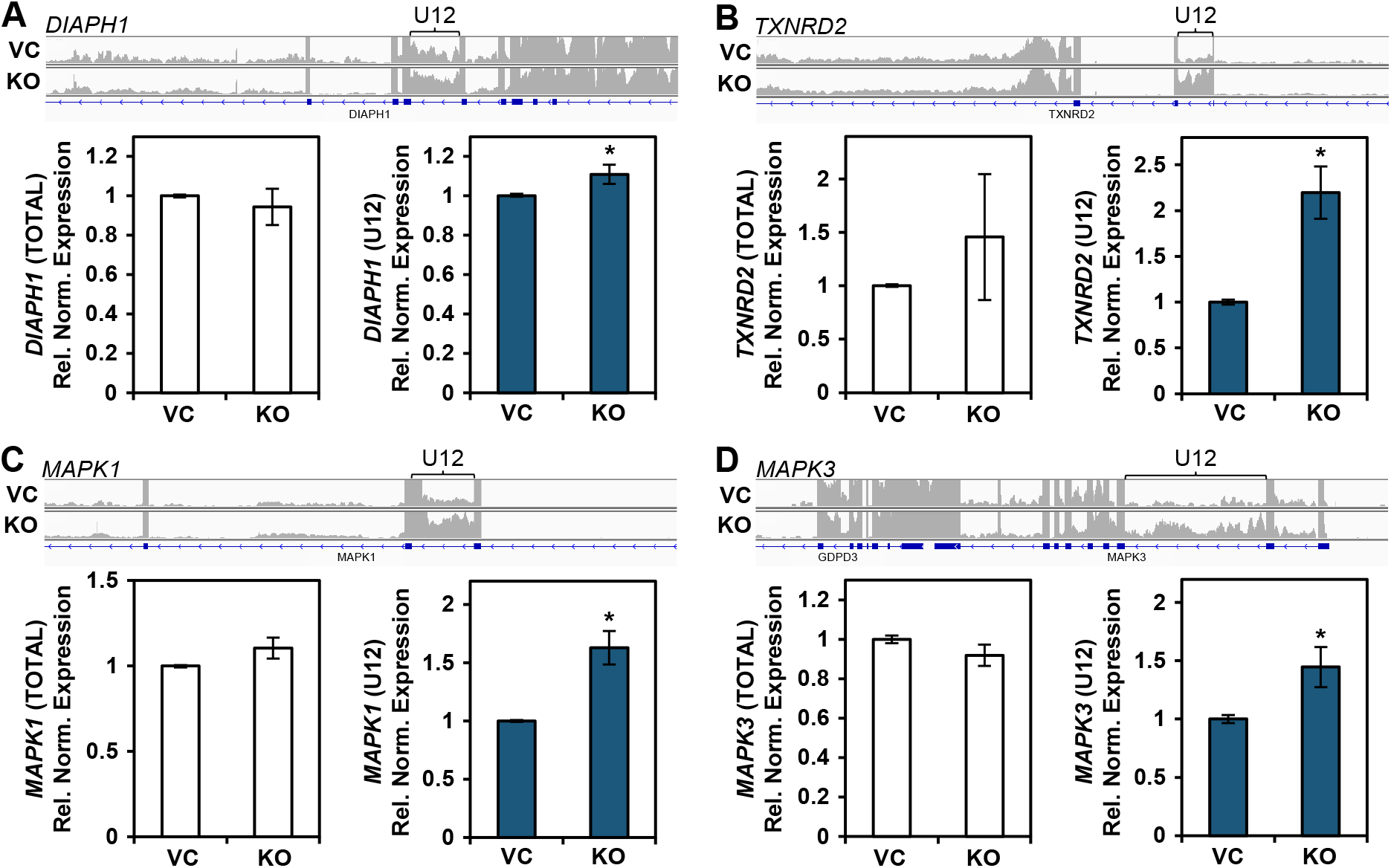
RT-qPCR validation of U12-type intron retention in RNA-seq libraries. **(A-D)** Summed read depth analysis from the six serial passages of VC and *RBM48* KO cells are shown at the top of each panel representing the regions of **(A)** *DIAPH1*, **(B)** *TXNRD2*, **(C)** *MAPK1*, and **(D)** *MAPK3* with significant (*q*<0.05) retention of a U12-type intron in KO cell populations compared to VC. Brace symbol indicates the U12-type intron with increased read depth in KO cells. The bottom panels display RT-qPCR analysis of the relative expression (mean ± SEM; n = 6 per subline) of each respective total transcript (left) and corresponding U12-type intron-containing transcript (right) from KO samples compared to VC. Expression was normalized to *HPRT1, IPO8*, and *PGK1* expression. Primers for RT-qPCR analysis are available in Table S5. \**p*<0.05.

Maize *rbm48* mutants cause specific retention of U12-type introns with impacts on U2-type introns that cannot be distinguished from random noise^25^. The log_2_ fold-change (log_2_FC) distributions between human *RBM48* KO and VC for all U2-type and U12-type introns are plotted in Figure 5A. As displayed, U12-type introns are shifted to greater overall log_2_FC values than U2-type introns. For significantly mis-spliced introns, in comparison to the observed 3.7% of U12-type introns retained in VC, genetic ablation of RBM48 drastically shifts intron retention events as 47.8% of U12-type introns are retained in KO (Fig. 5B). In contrast, only 9.1% and 8.6% of significant U2-type intron retention events occur in the KO and VC populations, respectively, with the minor enrichment observed in KO being well within the predicted noise from the expected 5% false positive threshold. These findings indicate that like maize^25^, loss of RBM48 in humans specifically increases retention or mis-splicing of U12-type intron sequences.

**Figure 5.**
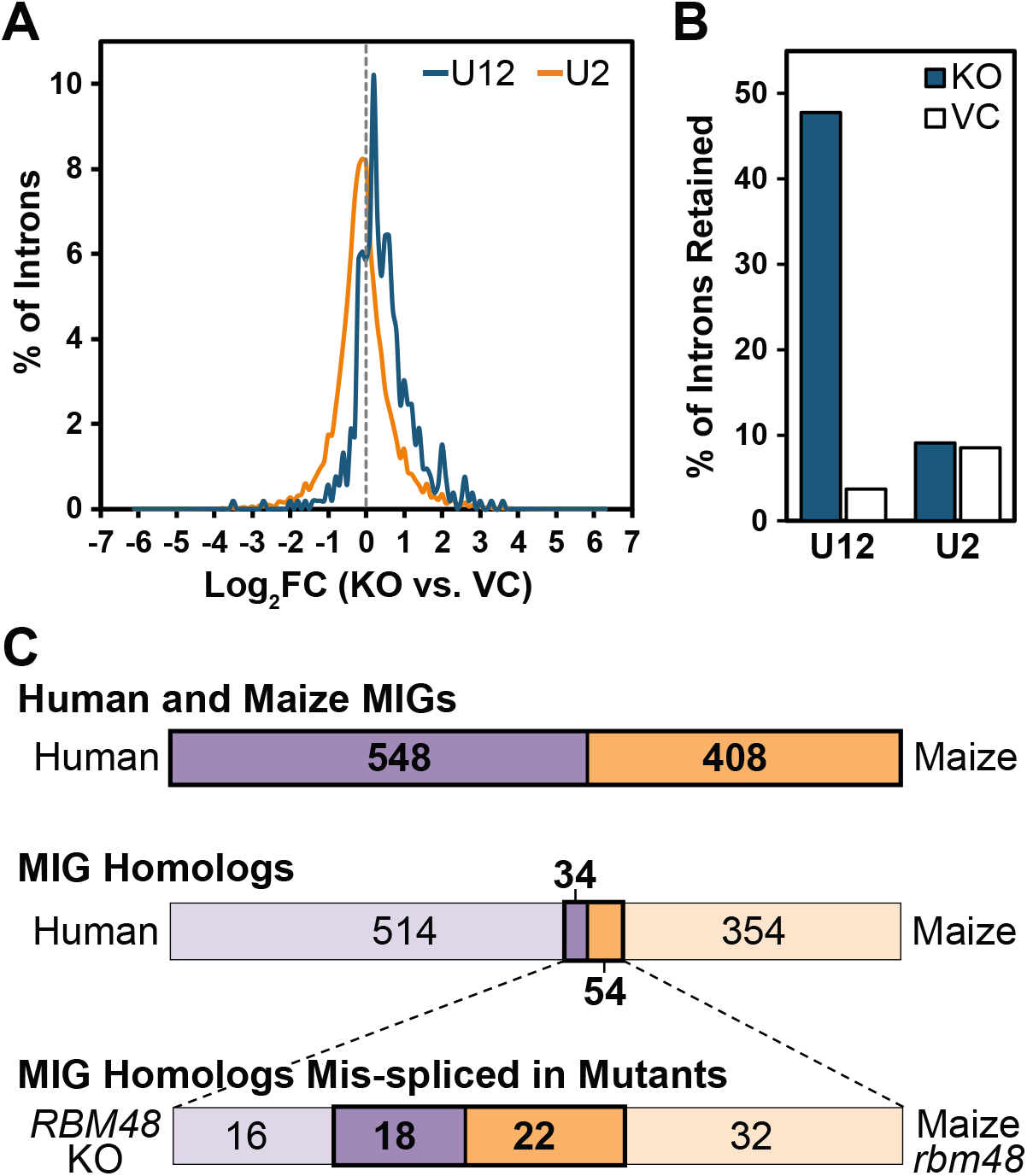
Comparative analysis of U12 intron retention between the RBM48 mutants of human and maize. **(A)** Log_2_ fold change (Log_2_FC) plot of intron expression in KO vs. VC samples. All introns sampled have a minimum sum of 10 intronic and exon-exon junction reads and an intronic read density >0 in both KO and VC samples. **(B)** Bar graph of the percent of introns significantly retained (q<0.05) in the KO vs. VC sample in U12-type and U2-type introns, respectively. **(C)** Schematic of overlapping *RBM48* splicing defects in homologous human and maize MIGs. Top bar shows total human (purple) and maize (orange) MIGs based on overlap between U12DB and MIDB (human) and ERISdb (maize). The middle bar shows the number of homologous MIGs between humans and maize based on reciprocal blastp analysis. Bottom bar shows homologous MIGs that are significantly mis-spliced in both human and maize *RBM48* mutants. Maize data is available from Bai *et al*. (2019)^25^.

### Conservation of mis-spliced genes shared between U12 splicing mutants of human and maize

The aberrant splicing of multiple U12-type introns poses a challenge to unambiguously identify target genes that cause defective phenotypes. We performed comparative analysis and searched for MIGs that display orthologous defects of U12-type intron splicing between the RBM48 mutants of human and maize. Gault *et al*. found 36 human MIGs that share homology with 57 maize MIGs^21^. With the higher confidence dataset for human U12-type introns used herein (see RNA-seq Methods), this homologous set is reduced to 34 human and 54 maize MIGs. These candidate orthologs have conserved domain structure and high sequence similarity of greater than 80-bit score using reciprocal blastp. The imbalance of human to maize MIGs is a result of differential gene duplication and divergent retention of duplicated genes since the split of animal and plant lineages^26^. Therefore, several single copy human MIGs share homology with two or more gene copies in maize. Of these 34 human MIGs, 21 (62%) are significantly mis-spliced in *RBM48* KO cells. As shown in Figure 5C, 18 human MIGs share significant mis-splicing with 22 homologous maize MIGs in *rbm48* endosperm (Table 1). To validate these RNA-seq comparisons, we selected four genes, *SMYD2, TAPT1, WDR91* and *ZPR1*, that showed significant U12-type intron retention in both human *RBM48* KO and maize *rbm48* mutants (Fig. 6). As shown in Figure 6A, the *RBM48* KO cells exhibited increased RNA-seq read depth in U12-type intronic regions compared to VC. When analyzed by RT-PCR, each of these genes also display stronger band intensity of the U12-type intron retained transcript in *RBM48* KO mutants (Fig. 6B). Moreover, corresponding RT-PCR assays of maize homologs show increased U12-type intron retention products in maize *rbm48* mutants relative to wild-type (Fig. 6B).

**Table 1.**
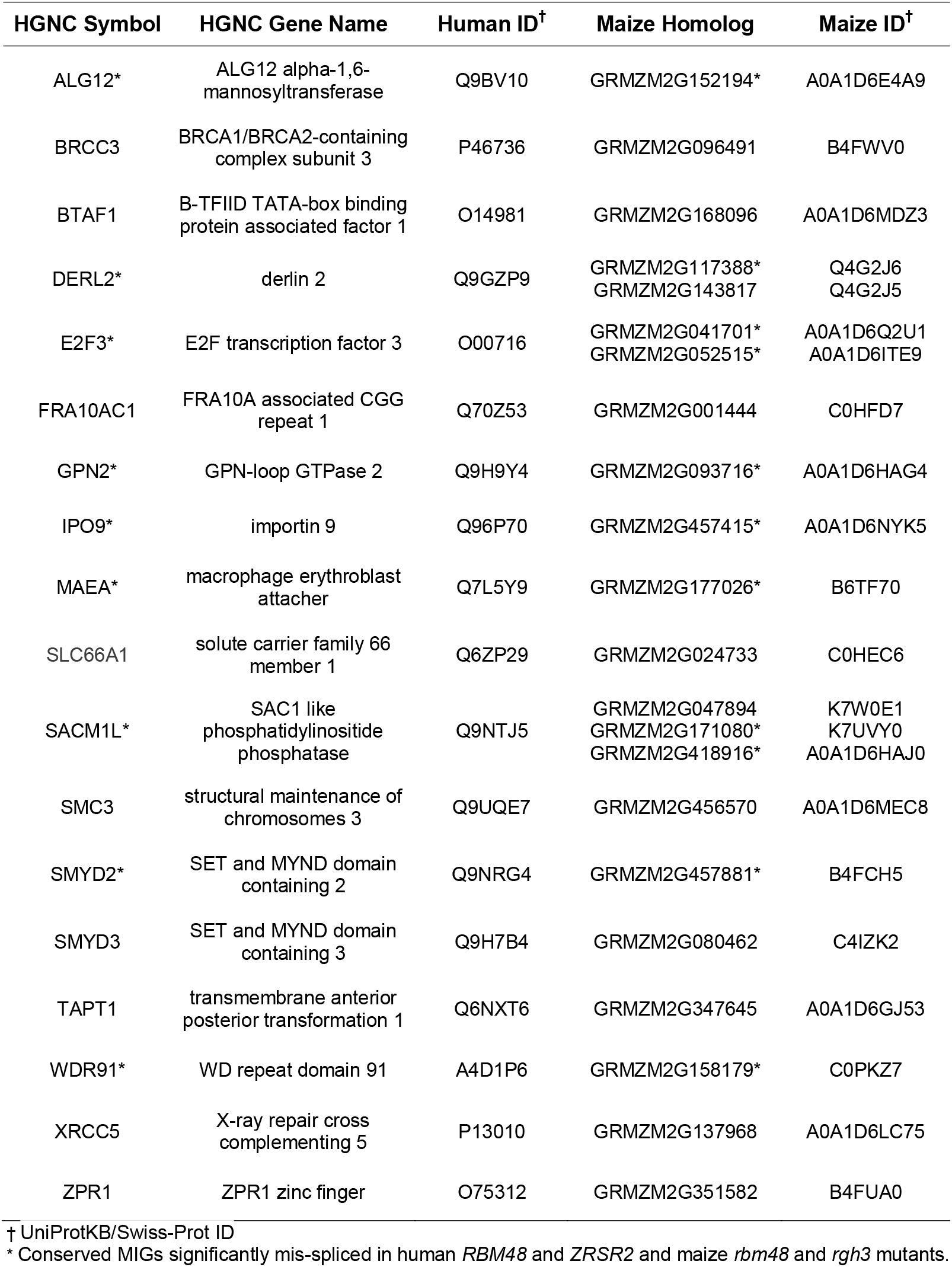
Conserved MIGs Mis-spliced in Human and Maize RBM48 Knockout Mutants.

**Figure 6.**
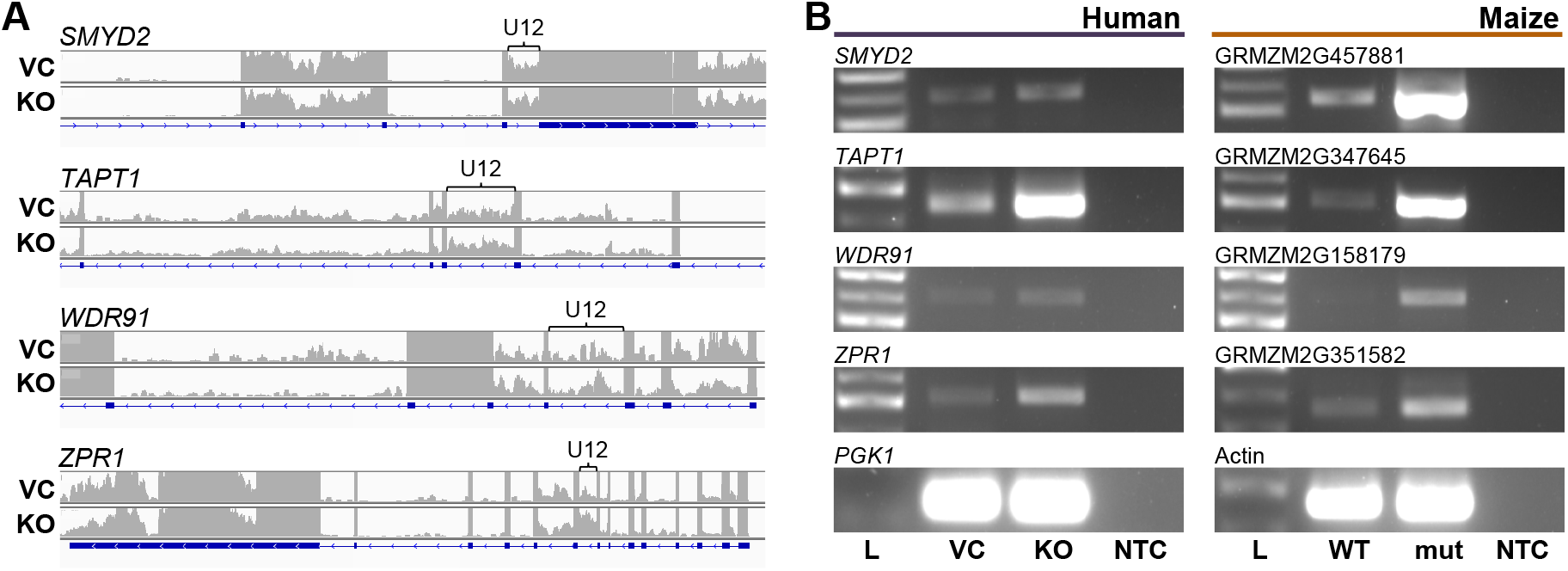
Comparison of U12-type intron retention in homologous MIGs between humans and maize. **(A)** Display of summed read depth analysis of significantly (*q*<0.05) increased U12-type intron retention in KO compared to VC cells for *SMYD2, TAPT1, WDR91*, and *ZPR1*. The brackets mark the position of the U12-type introns. **(B)** Semi-quantitative RT-PCR comparison of retention in U12-type introns indicated by band intensity in human (left panel) *SMYD2, TAPT1, WDR91*, and *ZPR1* and their maize (right panel) homologs GRMZM2G457881, GRMZM2G347645, GRMZM2G158179, and GRMZM2G351582, respectively. The position and sequence of the primers used during RT-PCR analysis are displayed in Figure S2 and Table S4, respectively.

A similar example of U12-type intron retention in conserved MIGs has been observed between human *ZRSR2* and maize *rgh3* mutant samples^21, 22^. Previously, we reported strong overlap of U12-type intron targets between maize *rbm48* and *rgh3* mutants as well^25^. Interestingly, of the 34 human and 54 maize homologous MIGs, 9 human MIGs are significantly mis-spliced in our *RBM48* KO and the *ZRSR2* mutants^22^ corresponding to 11 significantly mis-spliced homologous MIGs in the maize *rbm48*^25^ and *rgh3*^21^ mutants (Table 1).

## Discussion

Components of the minor spliceosome play important roles in promoting normal multicellular processes in both plants and animals. However, the mechanisms by which they impact cell development and differentiation are not well understood. Mutations in minor spliceosomal factors cause pleiotropic molecular defects affecting a large proportion of MIGs^15, 17, 21, 27-29^. This has made it difficult to delineate the causative genes underlying the pleiotropic phenotypes of MIG spliceosomal factor mutants, even if they exhibit highly similar characteristics.

Recently, cryo-electron microscopic analysis of purified human minor spliceosome structure revealed RBM48 and ARMC7 bind to the 5’ cap of U6atac^24^. In this context, we previously demonstrated the protein-protein interaction of RBM48 with ARMC7 is evolutionary conserved between maize and humans^25, 30^. Based on the role of maize RBM48 in U12-type intron splicing, we hypothesized human RBM48 would also impact U12-type intron splicing. We investigated RBM48 function by generating a CRISPR/Cas9 knockout in the human K-562 cell line and found that this role is conserved between humans and maize. Approximately 48% of U12-type introns analyzed in the *RBM48* KO subline were significantly retained, with their corresponding MIGs associated with several different pathways that are essential to growth and differentiation. We further determined that there are 18 orthologous MIGs mis-spliced in both human and maize RBM48 mutants. These observations identified a small group of MIGs as candidates for potential trans-species drivers of mutant phenotypes.

With the exception of *FRA10AC1*, mutations that either alter or abolish gene expression of these 18 conserved MIGs have been documented in the literature. In each case, the MIG mutations result in phenotypes displaying a broad spectrum of developmental defects in either humans or plants, several of which are reported in both species. For example, mutations in human ALG12 lead to Congenital Disorders of Glycosylation type Ig (CDG-Ig). CDG-Ig patients display a wide range of pleiotropic effects, including but not limited to hypotonia, developmental delay, and abnormal blood clotting^31-33^. A mutation in the homologous Arabidopsis ALG12 gene disrupts ER-mediated degradation of brassinolide receptors conferring a dwarf mutant phenotype to the plant^34^. Similarly, mutations in the chromatid cohesin *SMC3* and homologous *TTN* genes lead to pleotropic developmental defects in both humans and plants, respectively^35,36^.

Mutations in U12 splicing factors are also reported to inhibit cell differentiation and promote cell proliferation in both plants and animals. Somatic mutations of human *ZRSR2* cause suppression of myeloid cell differentiation and increased proliferation of myeloid progenitor cells^22^. Similarly, maize mutants in both *rgh3* and *rbm48* suppress endosperm differentiation and increase proliferation of endosperm in tissue culture^21, 25^. Within several of the conserved MIGs identified herein, mutations have been linked to abnormal cell proliferation or cancer in humans. For example, mutations in *BRCC3*, which encodes a cell cycle regulator involved in G2/M phase transition, have been linked to myelodysplastic syndrome (MDS) as well as breast and cervical cancers^37-39^. Additionally, mutations in *E2F3* have been associated with retinoblastoma, bladder and lung cancer^40-42^.

Despite these observed similarities, cell type- and lineage-specific differences play a large role in phenotypic effects of U12-type intron retention. Differential effects on *in vitro* cell viability are observed with targeted loss of *RBM48* in genome-wide CRISPR/Cas9 dropout screens where RBM48 is deemed essential in some human cell lines and dispensable in others^30, 43-47^. In a recent meta-analysis of 17 CRISPR screens conducted using three different large-scale libraries, *RBM48* is found to be essential for survival in 4 out of 7 leukemia-derived cell lines^44^. This includes the OCI-AML3, OCI-AML2, HL-60 and HAP1 lines. However, viability is unaffected in MV4-11, KBM-7 and K-562 cells deficient in *RBM48*, which agrees with our results in the *RBM48* KO K-562 subline. Interestingly, the genetic functions of *ZRSR2* or *RBM48* are highly overlapping in these 7 hematopoietic cell lines with *ZRSR2* and *RBM48* being essential in OCI-AML3, OCI-AML2, and HL-60, while both genes are dispensable in K-562, KBM-7, and MV4-11^44^. These 7 cell lines each retain a degree of lineage plasticity and can be induced to further differentiate in response to defined stimuli. For example, the adherent HAP1 cell line was isolated from an attempt to generate induced pluripotent stem cells from KBM-7 suspension cells and as a result, no longer express the hematopoietic markers CD43 and CD45^48^. These two nearly haploid lines also vary in their requirement of *RBM48* and *ZRSR2* for survival. These cell-type specific differences suggest that phenotypic effects of U12-type intron retention are likely dependent on the lineage-specific differentiation potential of each cell line with disruption of MIG-specific pathways resulting in lethality in some cell lines but not others.

The recent purified structure of the human minor spliceosome revealed six conserved amino acid residues of RBM48 that mediate the interaction of RBM48-ARMC7 protein complex with U6atac^24^. Of these, four amino acid residues are N-terminal to the conserved RRM domain of RBM48. Thus, the position of the sgRNA#1 is predicted to interrupt three of these residues, possibly negating a vital RBM48-U6atac interaction and rendering a lethal or anti-proliferative phenotype to the resultant host cells. In contrast, the position of sgRNA#2 spanning the distal region of the RRM domain does not impact the expression of these conserved residues, allowing targeted, mutant K-562 cell sublines to proliferate sufficiently for *RBM48* KO analysis. In vertebrates, RBM48 and ARMC7 are differentially compartmentalized in the nucleus and cytosol, respectively^49^ (Human Protein Atlas available from http://www.proteinatlas.org), which warrants investigation of *in vivo* significance of RBM48-ARMC7 interactions.

A combination of *in vivo* co-expression, Bimolecular Fluorescence Complementation (BiFC), and *in vitro* pull-down assays showed that maize RBM48 interacts with RGH3 and the core U2 splicing factors U2AF1 and U2AF2^24^. U2AF2 binds to the polypyrimidine tract of U2-type introns; whereas U2AF1 makes contact with the acceptor site^50^. Human ZRSR2 binds to the 3’ splice site of U12-type introns and is required for the formation of pre-spliceosomal complexes, but ZRSR2 also interacts with U2AF2 and is essential for the second transesterification reaction of U2 splicing^6^. These protein-protein interaction and *in vitro* splicing observations suggest that the major and minor spliceosomes interact for splice site selection. However, the splicing phenotypes for mutants in human *ZRSR2* as well as maize *rgh3* argue that *in vivo* protein functions are specific for U12-type introns^21, 22, 25^. The precise role of RBM48-ZRSR2 interactions in U12 splicing mechanisms remains to be determined. Perhaps, interactions of RBM48 with ZRSR2 may promote formation of the activated minor spliceosome.

Developmental gene expression is intricately regulated in a temporal, cellular, and tissue-specific manner beginning in the early stages of embryogenesis. Surprisingly, the inefficient splicing of U12-type introns often perturbs the coding potential but not the expression level of MIGs, many of which are essential for development. We found no changes in total transcript expression levels in our validation assays and similar observations have been made for maize *rgh3* and *rbm48*^21, 25^. Consequently, the mutant phenotypes observed for U12-dependent splicing defects are likely a direct result of the disrupted post-transcriptional processing of MIG transcripts. Further elucidation of the mechanism(s) by which retention of U12-type introns result in mutant phenotypes will not only entail biochemical and genetic analysis of individual MIGs but will also require comprehensive profiling of their expression in diverse cells and tissues at different stages of development, both at the levels of transcription and translation.

## Materials and Methods

### RBM48-Targeting CRISPR/Cas9 Constructs and Donor Templates

Vector and target-specific guide RNA (sgRNA) were designed as described^51^. The three *RBM48*-specific sgRNAs (Table S2) were separately cloned into the pX459 expression vector [pSpCas9(BB)-2A-Puro; Addgene plasmid ID: 48139]^52^. The single stranded DNA oligonucleotide for homology directed repair to generate a C-terminal *RBM48* Myc-epitope tag was obtained from Integrated DNA Technologies (IDT) (Table S2, Fig. S1).

### Transfection of K-562 Cells

K-562 cells (American Type Culture Collection (ATCC) CCL-243) were cultured in Iscove’s Modified Dulbecco’s Medium (IMDM) (Hyclone), supplemented with 10% (v/v) fetal bovine serum (FBS) (Hyclone) and incubated with 5% CO_2_ at 37°C. K-562 cells were seeded in triplicate into 6-well plates at a density of 2.4 × 10^5^ cells per well in 3 mL culture media for 24 h prior to transfection. For *RBM48* KO cells, each well was transfected with 0.5 μg pX459 construct expressing sgRNA#1 or sgRNA#2. VC cells were transfected with 0.5 μg pX459 alone. For *RBM48*-*Myc*-epitope tagging, cells were co-transfected with 0.5 μg pX459 construct with or without sgRNA TAG and 0.5 μg homology directed repair donor template. Vector DNA was mixed with 0.75 μL lipofectamine 3000 and diluted in Opti-Mem I media (Gibco/Thermo Fisher Scientific). Transfected cells were incubated for 24 h and then selected for 48 h using 3 μg/mL puromycin dihydrochloride. Transfectants were resuspended in fresh media and aliquots were isolated for cell viability assays and genotyping (day 0). Six-well culture plates were initially seeded with approximately 10^5^ cells per well in 3 mL culture media. Cell viability was monitored every 48 h for 16 days using trypan blue exclusion assay and counted by hemocytometer. Significant differences were determined with unpaired Student’s *t*-tests adjusted for multiple comparisons using the Benjamini and Hochberg False Discovery Rate. Viable cells from day 16 cultures of VC and sgRNA#2-targeted *RBM48* KO K-562 cells were subsequently cultured across six serial passages. Cells from each passage were collected and used for analyses representing six biological replicates of each subline. For the *RBM48*-*Myc* tagged co-transfectants, single-cell colonies isolated by serial dilution were expanded for two weeks prior to genotypic analysis. A single homogenous *RBM48-Myc* cell population was selected and similarly re-transfected with either pX459 or pX459 construct containing sgRNA#2. Following puromycin selection, cells from three serial passages of VC and heterogeneous sgRNA#2-targeted *RBM48-Myc* cultures were collected for analyses.

### Genomic DNA Extraction and Sanger Sequencing

Genomic DNA (gDNA) from harvested cell populations was extracted using the E.Z.N.A. Tissue DNA kit (Omega Bio-tek). DNA was amplified using *RBM48* specific primer pairs and amplicons were bidirectionally sequenced with the same or custom inner forward and reverse primers (Table S3). Sanger sequencing was performed by Genewiz. The sequencing chromatograms were analyzed visually and the frequency of indels generated at CRISPR target sites were quantified using Tracking Indels by Decomposition (TIDE) analysis^53^.

### Surveyor Nuclease Assay

The genomic region spanning the CRISPR target sites were PCR amplified using primers listed in Table S3. PCR products were hybridized to promote heteroduplex formation with the temperature cycle: 95°C for 10 min; 95°C to 85°C ramping at -2°C/s; 85°C to 25°C at - 0.3°C/s; 25°C for 1 min and 4°C hold. The resultant DNA products were subjected to SURVEYOR nuclease analysis following instructions provided by the manufacturer (IDT).

### RNA Extraction, cDNA Synthesis and RT-PCR

VC and *RBM48* KO cells from the six serial passages were pelleted and subsequently homogenized in TRIzol reagent followed by phase separation. Total RNA was extracted from the supernatants using the RNeasy Plus Universal Kit (Qiagen) per the manufacturer’s protocol with the inclusion of gDNA digestion using the RNase-Free DNase Set. gDNA-free total RNA was eluted in 30 µL RNase free water and the RNA concentration and purity was determined by Nanodrop 2000C. RNA integrity was confirmed by gel electrophoresis using 1% agarose with ethidium bromide. gDNA-free total RNA (1 μg) was reverse transcribed using SuperScript VILO cDNA Synthesis Kit (Life Technologies). Reverse transcription (RT) was performed for 10 min at 25°C, 60 min at 42°C, and reactions were terminated by incubation at 85°C for 5 min. RT samples were used immediately or stored at -20°C. After first strand cDNA synthesis, MIG transcripts were amplified using gene-specific primers as shown in Figure S2 and listed in Table S4.

### RT-qPCR

RT-qPCR was performed in the Bio-Rad CFX96 Real Time System with target genes summarized in Table S5. Reaction mixtures consisted of 1 ng cDNA template, 400 nM specific sense primer, 400 nM specific antisense primer, RNase/DNase-free water, and 1x SsoAdvanced SYBR Green Supermix (Bio-Rad) in a final volume of 10 μL. The thermal profile of the PCR followed the SsoAdvanced supermix protocol: initial denaturation at 95°C for 30 s followed by 40 cycles of 95°C for 5 s and 60°C for 20 s with amplification data collected at the end of each cycle. Product specificity was validated with dissociation curves by incubating reactions from 65 to 95°C in 0.5°C increments for 5 s each. The Bio-Rad PrimePCR RNA quality, DNA contamination control, and Positive PCR control assays evaluated RNA quality, genomic DNA contamination, and PCR reaction performance respectively. cDNA from each of the six serial passages of the VC and *RBM48* KO cell populations were amplified in triplicate for each target gene.

PCR priming efficiencies of U12-type intron retained RNA templates were determined with calibration curves from the K-562 *RBM48* KO subline. cDNA (5 ng/50 μL reaction) was pre-amplified for 10 cycles with U12-type intron specific primers (Table S5). PCR amplicons were purified and quantified by NanoDrop 2000C. Input DNA copy number was determined using the University of Rhode Island Genomics and Sequencing Center online calculator (https://cels.uri.edu/gsc/cndna.html) with templates ranging from 2 × 10^3^ to 2 × 10^6^ copies/well. Alternatively, a 10-fold dilution series of cDNA from K-562 ATCC CCL-243 cells ranging from 5 pg to 50 ng cDNA/well was assayed. The quantification cycle (Cq) was plotted against the log amount of cDNA input and the relationship between Cq values and RNA concentration was calculated by linear regression to find a slope and intercept that predicts cDNA amounts and correlation coefficient (R^2^). Amplification efficiencies (E) were calculated according to the equation E=(10(−1/slope)-1) × 100 and are expressed as a percentage. The qPCR parameters providing the standard curve for each primer pair are summarized in Table S6. A cDNA positive control inter-run calibrator (1 ng/well) was included in every run and the ΔCqs were verified to be <0.01 between runs. Six reference genes were included and validated with RefFinder^54^. The three genes with the lowest geometric mean rank having a geNorm^55^ stability value <0.5 were selected. Gene expression was determined using the Bio-Rad CFX Manager software version 3.0 and calculated using the efficiency corrected model^56^ of the ΔΔC_q_ method^57^ modified for normalization by geometric averaging of multiple reference genes^55^. Results are expressed as the ratio of *RBM48* KO ΔCq expression to VC ΔCq expression (Relative Normalized Expression). Significant differences were identified with unpaired Student’s *t*-tests.

### Analysis of RBM48-Myc Protein Expression

Total protein was extracted from VC and sgRNA#2-targeted *RBM48-Myc* cell pellets homogenized in RIPA lysis buffer (50 mM Tris-HCl, 150 mM NaCl, 1.0% Triton X-100, 0.25% sodium deoxycholate, 5.0 mM EDTA) in the presence of 10 μL/mL HALT protease inhibitor cocktail (Thermo Fisher Scientific). Extracts were centrifuged at 15,000 × g for 15 min at 4°C. Supernatant aliquots were frozen in liquid nitrogen and stored at -80°C. Total proteins were separated with 7.5% Mini-PROTEAN TGX sodium dodecyl sulfate polyacrylamide gel (Bio-Rad) with 20 µg of protein per lane, quantified via Bradford assay. Protein gels were wet transferred to PVDF membrane (Millipore). Specific proteins were detected using 1:1000 dilution of Myc-Tag (9B11) mouse monoclonal and proliferating cell nuclear antigen (PCNA, D3H8P) XP Rabbit monoclonal primary antibodies (Cell Signaling Technology). Following primary antibody incubation, membranes were washed and incubated with 1:2000 dilution of horseradish peroxidase-conjugated horse anti-mouse IgG (Cell Signaling Technology) and goat anti-rabbit IgG (Santa Cruz Biotechnology) secondary antibodies. Myc-Tag or PCNA signal was detected using SuperSignal West Femto (Thermo Fisher Scientific) or Clarity (Bio-Rad) ECL substrates, and imaged using the Bio-Rad ChemiDoc Touch imaging system. Chemiluminescent signals were quantified with the Image Studio Lite program version 3.1 (LI-COR Biosciences) to normalize the RBM48-Myc protein band density to the PCNA loading control. Statistical analyses were performed using unpaired Student’s *t*-tests.

### RNA-seq

Approximately 10^6^ cells were collected from each of six serial passages for both VC and *RBM48* KO sublines. These 12 biological samples were homogenized in TRIzol and subjected to chloroform/isopropanol extraction per the manufacturer’s instructions. Following DNase treatment, the samples were purified using RNeasy MinElute Cleanup Kit (Qiagen). RNA quality was confirmed to have an RNA Integrity Number (RIN) of at least 9.5 with the Agilent Tape-station 2200. Library construction and sequencing were completed at the University of Florida Interdisciplinary Center for Research. The NEBNext Ultra RNA Library Prep Kit for Illumina (New England Biolabs) was used to generate cDNA libraries. The libraries were then pooled and 2 lanes of Illumina HiSeq3000 paired-end 100 bp sequences were generated with a total of 750,648,520 reads with individual samples ranging 56,844,183 – 75,114,017 reads and averaging 62,554,043 reads.

Raw RNA-seq data were screened to remove adapter sequences using Cutadapt v1.1^58^ with the following parameters: --error-rate=0.1 --times=1 --overlap=5 --minimum-length=0 -- adapter=GATCGGAAGAGCACACGTCT --quality-base=33. Adapter trimmed sequences were quality filtered/trimmed with Trimmomatic v0.22^59^ using parameters (HEADCROP:0, LEADING:3, TRAILING:3, SLIDINGWINDOW:4:15, and MINLEN:40) to truncate reads for base quality <15 within 4 base windows and kept only reads ≥40 bases after trimming. Only reads remaining in pairs after Trimmomatic were used for subsequent analysis. On average, 28,839,533 read pairs per sample (range 26,349,516-33,018,146) remained after quality filtering.

Reads were aligned to the human genome sequence assembly (GRCh38) with HiSAT2^60, 61^ using the following parameters: --max-intronlen 100000 -q --pen-noncansplice 6 -- no-discordant --rna-strandness RF. Homo_sapiens.GRCh38.87. Annotation (gtf/gff) was used for intron counts and transcripts per million (TPM) normalization. A custom GFF file detailing all non-redundant introns from Homo_sapiens.GRCh38/hg38 was constructed and used to perform U12-type intron retention analysis. Introns within this GFF file were given a unique identifier and annotated as U12- or U2-type introns. Determination of intron type (U12 or U2) was first based on a publicly available collection of human U12-type introns^9^ available at https://www.crg.eu/en/programmes-groups/guigo-lab/datasets/u12db-database-orthologous-u12-type-spliceosomal-introns. Six-hundred and ninety-five U12-type introns were identified by Alioto^9^ and their sequences and coordinates relative to Human genome annotation version NCBI35/hg17 were available. Conversion of the NCBI35/hg17 iU12-type intron coordinates to Homo_sapiens GRCh38/gh38 was accomplished using a two-step process. The annotation “lift-over” utility available from the UCSC genome browser (https://genome.ucsc.edu/cgi-bin/hgLiftOver) was used to first convert the U12-type intron coordinates from NCBI35/hg17 to CRCh37/hg19, and then again from CRCh37/hg19 to Homo_sapiens.GRCh38.87. Given the age of the annotation data contained within U12DB, this list was then cross-referenced to the recently available collection of human U12-type introns within the minor intron database (MIDB)^62^ available at https://midb.pnb.uconn.edu. To generate a high confidence list, all introns that did not overlap between the two databases were removed from our dataset.

Read counts/gene were determined with the HTSeq-Count utility in the HTSeq package (Ver 0.8.0)^63^ using the start-stop coordinates of the entire locus and the following parameters: -m intersection-nonempty -s reverse). Intron counts were determined with the HTSeq-Count utility (-m intersection-nonempty -s reverse) using the intron GFF file for features. Significant differences in summed intron and exon-exon junction read counts between KO and VC populations were determined by Fisher’s Exact Test. Multiple comparison adjustment was performed using the Benjamini-Hochberg False Discovery Rate (FDR) and reported as the adjusted q-value with a significance threshold of 0.05.

Gene ontology (GO) term enrichment analyses used String-DB with default parameters and the genes having greater than 0 TPM in either VC or KO (genes expressed in either K-562 cell subline) as a customized reference. MIGs analyzed were selected based on significant retention (*q*≤0.05) of U12-type introns in *RBM48* KO. Non-coding MIGs were excluded. String-DB results were further filtered by enrichment cutoff of >2.0 and *q*<0.01. GO term redundancy was filtered with ReviGO using Simrel semantic similarity measurement with allowed similarity=0.5.

VC and KO population intron reads and their respective flanking exon-exon junction reads were first filtered for a minimum sum of 10 intronic and exon-exon junction reads in both VC and KO. Read densities were then calculated for intron density (Di) and exon-exon junction density (De) separately as described^64^. Introns were selected for analysis when Di>0 to calculate log_2_ fold-change of intron splicing efficiency between *RBM48* KO and VC.

### Analysis of MIG Homology

Putative orthologs of maize and human MIGs were identified with reciprocal blastp searches against human and maize protein sequence databases. Human and maize MIGs that returned reciprocal hits with bit scores >80 were considered orthologous. We searched for homologous MIGs that share significant intron retention between human *RBM48* KO population and maize mutant *rbm48*^25^. The position of the intron was determined by direct splice alignment of the protein sequence by SplicePredictor^65^. The position of the intron was considered conserved if the position was within 5 amino acids residues between the two species.

## Acknowledgments

This work was supported by the National Science Foundation (grant 1412218 to S.L., W.B.B. and A.M.S.); the National Heart Lung and Blood Institute at the National Institutes of Health (grant R01-HL135035 to R.J.W.); the National Cancer Institute at the National Institutes of Health (grant R15-CA182889 to G.J.M.); the Oakland University Research Excellence Fund (to S.L., G.J.M. and R.J.W.) and American Heart Association Innovative Research Grant (to R.J.W.).

## Author Contributions

A.E.S., G.J.M., and S.L. designed research; A.E.S., J.C., J.P.G., L.L., L.M.H., C.K., R.D., and W.B.B. performed research; A.E.S., J.C., J.P.G., W.B.B., R.J.W., G.J.M., and S.L. analyzed data; and A.E.S., J.C., A.M.S., R.J.W., and S.L. wrote the paper.

