## Supplemental Figures and Tables for "Genetic Analysis of Human RNA Binding Motif Protein 48 (RBM48) Reveals an Essential Role in U12-Type Intron Splicing"

**This PDF file includes:**

Figures S1 to S2

Tables S1 to S6

Dataset S1 Legend

Dataset S2 Legend

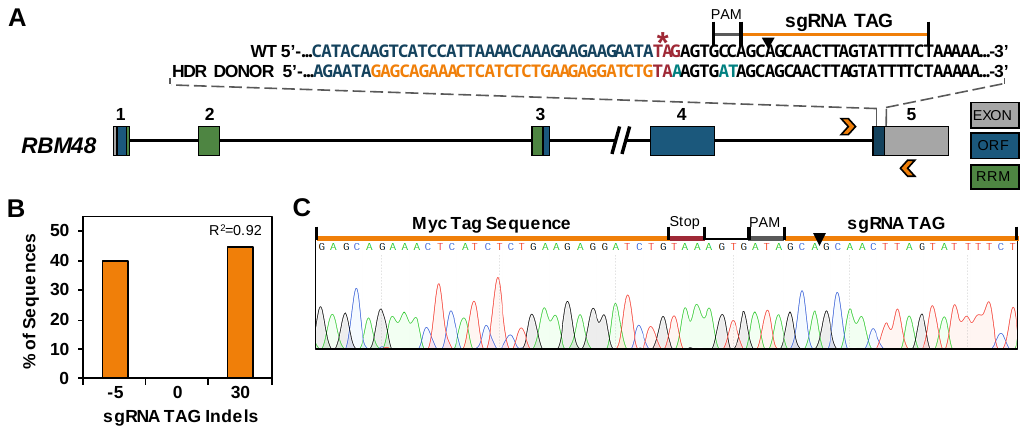

Fig. S1. CRISPR/Cas9 mediated C-terminal epitope tagging of *RBM48* in K-562 cells.(A) Schematic of *RBM48* (as described in Figure 1) displaying the design and position of the sgRNA TAG used for Cas9 targeting of the C-terminal region of *RBM48* and the ssDNA donor template utilized for homology directed repair (HDR). The wild-type (WT) and HDR Donor sequences of the modified region are shown. The HDR donor template consists of a sense 193 bp sequence spanning genomic coordinates 92536858-92537020 of chromosome 5 (GRCh38.p4) and encodes the 30 bp Myc-tag (orange text) flanking the stop codon of *RBM48* (red text and asterisk). To prevent donor template folding and misincorporation into the genome, a G to A transversion mutation within the *RBM48* stop codon and two mutations within the PAM that also serve to prevent donor DNA cleavage by Cas9 were included (teal text). In this design, 77 bp homology arms flank the 30 bp insert and the 3’-most PAM mutation with the position of the double-strand break occurring 4 bp upstream of the 3’ homology arm junction. Arrowheads indicate position of RBM48 Tag_Out primers (Table S3; nested RBM48 Tag_In primers not shown) utilized for analysis in (B) and (C). (B) TIDE analysis of *RBM48-Myc* K-562 gDNA derived from the isolated cell colony utilized in these studies. The analysis reveals mono-allelic incorporation of the Myc epitope tag into the *RBM48* genomic locus (indicated by the 30 bp insertion) with the remaining allele containing a 5 bp deletion within the 3’-UTR of *RBM48* (sequence data not shown). (C) Sanger sequencing chromatogram of the TOPO-TA cloned *RBM48* Myc-tagged allele displaying correct sequence incorporation of the HDR Donor template into the K-562 cell genome.

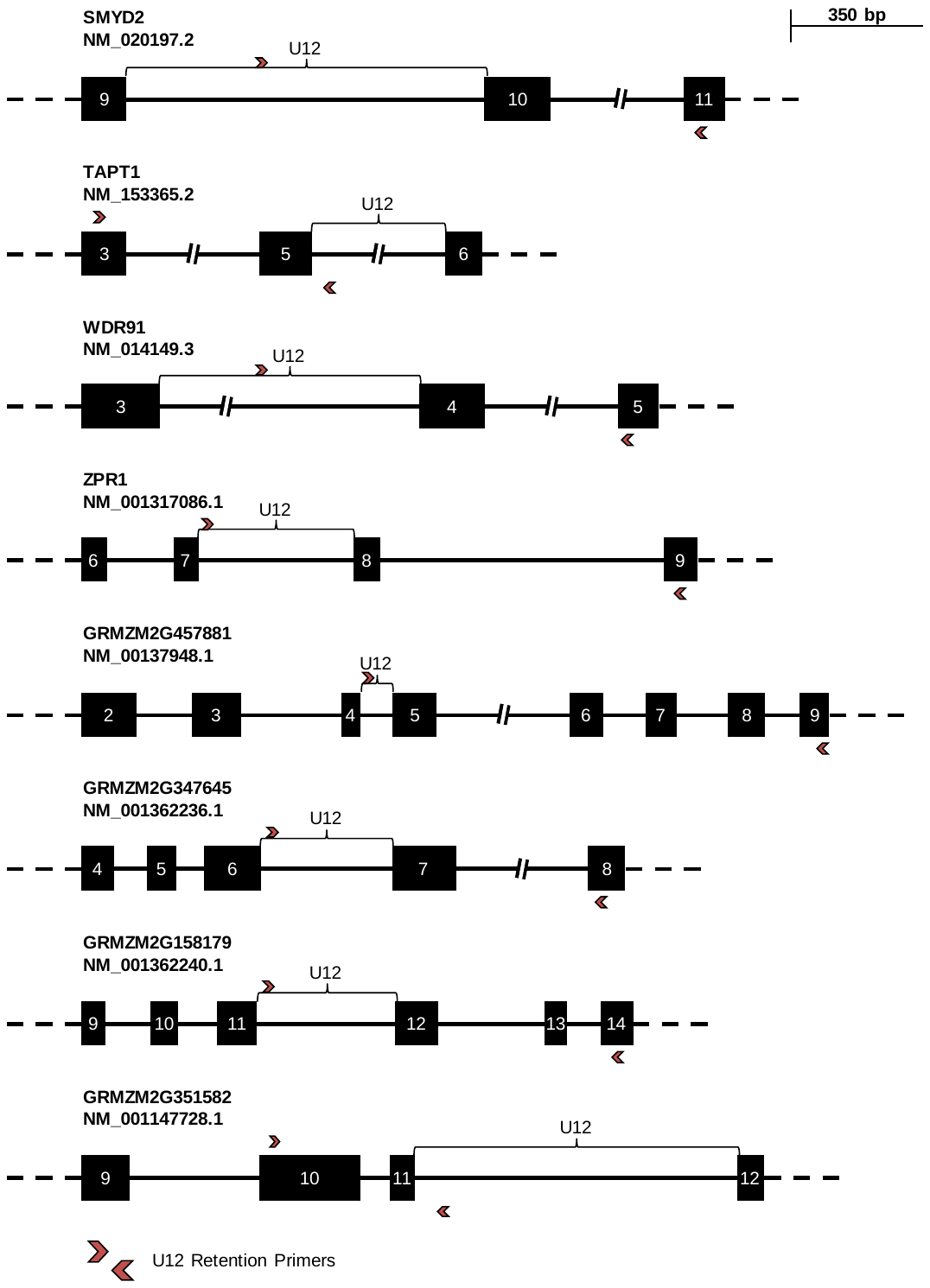

Fig. S2. Schematic of primer positions used in comparative RT-PCR analysis of homologous MIGs shared between humans and maize. Exons and introns are represented as boxes and lines, respectively. Dotted lines indicate the flanking transcript sequence. Brackets mark the position of U12 introns. Red arrowheads mark the position of the primers used to detect U12 intron retention. The primer sequences are listed in Table S4.

Table S1. Enriched Biological Processes among significantly retained MIGs

| **GO Term** | **Description** | **Retained MIGs** | **Expressed Coding Genes** | **FDR** |
| --- | --- | --- | --- | --- |
| GO:0051641 | cellular localization | 85 | 2115 | 2.76E-11 |
| GO:0033036 | macromolecule localization | 78 | 2201 | 4.09E-08 |
| GO:0071705 | nitrogen compound transport | 60 | 1636 | 1.56E-06 |
| GO:0007049 | cell cycle | 48 | 1234 | 1.15E-05 |
| GO:0048193 | Golgi vesicle transport | 20 | 317 | 0.00014 |
| GO:0033554 | cellular response to stress | 51 | 1505 | 0.0002 |
| GO:0044403 | symbiont process | 28 | 627 | 0.0005 |
| GO:0018206 | peptidyl-methionine modification | 5 | 11 | 0.00056 |
| GO:0042147 | retrograde transport, endosome to Golgi | 9 | 78 | 0.0012 |
| GO:0009894 | regulation of catabolic process | 32 | 824 | 0.0012 |
| GO:0017196 | N-terminal peptidyl-methionine acetylation | 4 | 6 | 0.0013 |
| GO:0016482 | cytosolic transport | 11 | 131 | 0.0019 |
| GO:0044770 | cell cycle phase transition | 15 | 258 | 0.0036 |
| GO:0006974 | cellular response to DNA damage stimulus | 28 | 728 | 0.0037 |
| GO:0043647 | inositol phosphate metabolic process | 7 | 55 | 0.0046 |
| GO:0051276 | chromosome organization | 33 | 959 | 0.0054 |
| GO:0031365 | N-terminal protein amino acid modification | 5 | 25 | 0.0067 |
| GO:0036503 | ERAD pathway | 8 | 84 | 0.007 |
| GO:0070925 | organelle assembly | 25 | 652 | 0.007 |
| GO:0006913 | nucleocytoplasmic transport | 14 | 254 | 0.0072 |
| GO:0070201 | regulation of establishment of protein localization | 24 | 629 | 0.0088 |
| GO:0006259 | DNA metabolic process | 27 | 748 | 0.0088 |
| GO:0071276 | cellular response to cadmium ion | 5 | 28 | 0.0088 |
| GO:0019886 | antigen processing/presentation of exogenous peptide antigen via MHC class II | 8 | 89 | 0.0088 |

Table S2. Sequences of Oligonucleotides Used as HDR Donor Templates and for Cloning sgRNA

| **Template** | **Sequence 1** | **Sequence 2** |
| --- | --- | --- |
| sgRNA#1 | CACCGTCAGGTATATACAATCAATT | CAGTCCATATATGTTAGTTAACAAA |
| sgRNA#2 | CACCGCTAGAAAAAAACTACAAATG | CGATCTTTTTTTGATGTTTACCAAA |
| TAG sgRNA | CACCGAGAAAATACTAAGTTGCTGC | AAACGCAGCAACTTAGTATTTTCTC |
| TAG sgRNA  HDR Donor | CATCTGTGCCAAAGCCTCCAGAGGACAAGCCAGAAGATGTACATACAAGTCATCCATTAAAACAAAGAAGAAGAATA**GAGCAGAAACTCATCTCTGAAGAGGATCTG**TAAAGTGATAGCAGCAACTTAGTATTTTCTAAAAAGAACATTTATTATTTATTTTTAGCCTGTCATTTTAATTC TTCAAGAGATTT | |

Table S3. Genotyping Primers

| **Primer Name** | **Primer 1 Sequence** | **Primer 2 Sequence** |
| --- | --- | --- |
| RBM48#1 | TTCCCAGTGACTTCTACCGA | GGCTGGAGGAAGATATGCTAGATT |
| RBM48#2 | TGTCCACAAGCAGAGCATCTT | AAACATCTTGCCTGGCTTGC |
| RBM48 Tag_Out | GCATATGTTCACTTTTCTTCCTCCA | GACGCTGGCTGCCTATCTTTATT |
| RBM48 Tag_In | AGGCTGTACCTTGAACTTAGGC | AGTTTCTGCAACATTAAGTATGGGT |

Table S4. RT-PCR Primer Sequences

| **Primer Name** | **Primer 1 Sequence** | **Primer 2 Sequence** |
| --- | --- | --- |
| *SMYD2* | GAAAGGCACATTGTTTCTCAGC | CCCTAGCTTCAACCACATGGA |
| *TAPT1* | GTGCCTGGATGCGTTTTTGT | GGTAGACTATGCATTAACTGTCGG |
| *WDR91* | CTCAGTCCAAACCTTTGCCA | TCAGTCGGTGGATTTCAGCTT |
| *ZPR1* | TCTTTTGCCAAGCAGTTGGG | ATGTGGAGGGTGATCCTGGT |
| *PGK1* | GTAAAGTCCTTCCTGGGGTGG | TAGCTAATGCCAAGTGGAGATGC |
| GRMZM2G457881 | CTCGTATCCTGCCGTGTTCA | GGCAGCAAAAAGCCATCACA |
| GRMZM2G347645 | TCTTGCTAGTCACTGTAGGTTAAGT | GGCCATGATGTGAAATCGCT |
| GRMZM2G158179 | CATTGTGATATTCTTGTACCATCCC | AGCAGCCTCTTGCCATTTGA |
| GRMZM2G351582 | TCGTGAACAACAAGCAGCAC | TGGCAACATAGGTGTTGTGAGT |
| Actin | CATGAGGCCACGTACAACTCCATC | TCATACTCTCCCTTGGAGATCCAC |

Table S5. RT-qPCR Primers

| **Gene** | **RefSeq Accession** | **Primer 1 Sequence**  **Primer 2 Sequence** | **Primer Location** | **Product Size (bp)** |
| --- | --- | --- | --- | --- |
| *RBM48* | NM_032120.4 | TCATCTGTGCCAAAGCCTCC  GCTGGCACTCTATATTCTTCTTCT | Exon 5  Exon 5 | 90 |
| *DIAPH1* (Total) | NM_001079812.3 | TACGATAGCCGGAACAAGCA  GGCTCTGACCAGCAGTAGGA | Exon 6  Exon 7 | 117 |
| *DIAPH1* (U12) | NM_001079812.3 | TGCAGGACCTTCGAGAGATTG  CATTCCACACAGGGATCAGGG | Exon 9/10  Intron 10 | 199 |
| *MAPK1* (Total) | NM_002745.5 | CAAGGGCTACACCAAGTCCA  GGTCAAGATAATGCTTCCCTGG | Exon 4/5  Exon 5 | 101 |
| *MAPK1* (U12) | NM_002745.5 | AGCACCAACCATCGAGCAAA  TTACCAAGCAGTGGAATTGGC | Exon 2  Intron 2 | 71 |
| *MAPK3* (Total) | NM_001109891.1 | ACATCTCTCATGGCTTCCAGG  GGCCATCAAGAAGATCAGCC | Exon 2  Exon 2 | 150 |
| *MAPK3* (U12) | NM_001109891.1 | ACCCTGGAAGCCATGAGAGA  ACAGAAACCAAGCAACGGGT | Exon 2  Intron 2 | 100 |
| *TXNRD2* (Total) | NM_006440.5 | ATCATTGCTACTGGAGGGCG  TTTCCAGGGGATTCCTTCAGC | Exon 7  Exon 8 | 107 |
| *TXNRD2* (U12) | NM_006440.5 | AGGTGCCTTGGAATATGGAATCA  AGGGAAGGAGTGTCCAGTTC | Exon 8  Intron 9 | 140 |
| *HMBS* | NM_000190.4 | AGAGAAAGTTCCCGCATCTGG  GTTGTGCCAGCCCATGC | Exon 8  Exon 9 | 137 |
| *HPRT1* | NM_000194.3 | GCTTTCCTTGGTCAGGCAGT  GGCTTATATCCAACACTTCGTGG | Exon 6  Exon 7 | 90 |
| *IPO8* | NM_006390.4 | TGCACGTCTCAGGTTTTTGC  TCATGTACAACAGAAGGCACTGT | Exon 25  Exon 25 | 72 |
| *PGK1* | NM_000291.4 | GTAAAGTCCTTCCTGGGGTGG  TAGCTAATGCCAAGTGGAGATGC | Exon 11  Exon11 | 120 |
| *TBP* | NM_003194.5 | TCCACAGTGAATCTTGGTTGTA  GGTTCGTGGCTCTCTTATCCTC | Exon 4  Exon 5 | 120 |
| *YWHAZ* | NM_003406.4 | AGATTCTGAACTCCCCAGAGAAAG  TCAGCTTCGTCTCCTTGGGTA | Exon 4  Exon 6 | 175 |

Table S6. Real Time PCR Efficiencies of Validated Reference Genes and Genes of Interest

| **Gene** | **Slope** | **Intercept** | **R^2^** | **Efficiency** | **Dilution Range** |
| --- | --- | --- | --- | --- | --- |
| *RBM48* | -3.312 | 41.98 | 0.991 | 100.4% | 5 pg – 50 ng |
| *DIAPH1* (Total) | -3.299 | 36.97 | 0.999 | 100.9% | 5 pg – 5 ng |
| *DIAPH1* (U12) | -3.465 | 30.54 | 0.996 | 94.4% | 2x10^3^ – 2x10^6^ copies |
| *MAPK1* (Total) | -3.332 | 35.41 | 0.998 | 99.6% | 5 pg – 5 ng |
| *MAPK1* (U12) | -3.345 | 32.11 | 0.995 | 99.0% | 2x10^3^ – 2x10^6^ copies |
| *MAPK3* (Total) | -3.354 | 38.38 | 0.998 | 98.7% | 5 pg – 5 ng |
| *MAPK3* (U12) | -3.428 | 31.57 | 0.999 | 95.8% | 2x10^3^ – 2x10^6^ copies |
| *TXNRD2* (Total) | -3.228 | 38.05 | 0.999 | 104.1% | 5 pg – 5 ng |
| *TXNRD2* (U12) | -3.445 | 30.95 | 0.998 | 95.1% | 2x10^3^ – 2x10^6^ copies |
| *HPRT1* | -3.412 | 38.3 | 0.993 | 96.4% | 50 pg – 50 ng |
| *IPO8* | −3.350 | 39.94 | 0.990 | 98.8% | 50 pg – 50 ng |
| *PGK1* | -3.328 | 37.32 | 0.994 | 99.8% | 5 pg – 50 ng |

Dataset S1 (separate file). Read counts and statistics for individual introns and exon-exon junctions detected in RNA-seq experiments. Intron coordinates are given for the Homo_sapiens.GRCh38.87 reference genome annotation. Intron reads and exon-exon junction reads are summed by Vector Control (VC) and Knockout (KO) K-562 sublines for all biological replicates. Introns were filtered for overlap with the MIDB dataset (see RNA-seq Methods in main article). Prior to analysis, introns were filtered for a minimum sum of 10 total exon-exon junction and intronic read counts with an intronic density (Di) > 0 in both VC or KO populations.

Dataset S2 (separate file). Read counts for individual introns and exon-exon junctions for each biological replicate from Vector Control (VC) and Knockout (KO) K-562 sublines detected in RNA-seq experiments.
